# How Much Does Protein Structure Really Help? A Case Study in Mutation-Induced Stability Prediction

**DOI:** 10.64898/2025.12.22.694225

**Authors:** Asher Moldwin, Amarda Shehu

## Abstract

Multimodal neural networks integrating protein language models (PLMs) with structure-derived features are increasingly common for predicting mutation effects, yet fundamental mechanisms remain poorly characterized. In this paper, we articulate and address two key questions: (i) do these architectures exploit mutation-conditioned structural changes, and (ii) does structure provide additive value over PLM-learned sequence embeddings afterall? Using Δ*T_m_* (melting temperature shift) prediction as a controlled testbed, a phenotype expected to depend strongly on three-dimensional geometry, we introduce generalizable diagnostic methodologies: systematic channel ablations quantifying each modality’s marginal contribution, and *context-radius probing*, a novel technique restricting inputs to progressively larger neighborhoods around mutations to spatially localize predictive signal. Across ten independent runs per condition, we find PLM embeddings dominate: removing structure causes minimal performance change, while removing PLMs causes performance collapse. Context-radius probing reveals signal is highly localized; mutation-site-only models recover full-context performance. Critically, comparing wild-type-shared versus mutation-conditioned structural regimes reveals no systematic gain from geometric perturbations, demonstrating that current representations function as static fold priors because downstream featurization attenuates mutation-induced changes. However, structure helps selectively: benefits concentrate in variants with atypical PLM embeddings occupying phenotypically incoherent neighborhoods where sequence-derived priors are locally unreliable. Though focused on a controlled testbed, this work surfaces a key challenge for any protein prediction task where domain knowledge suggests structure should matter: not whether to “add structure,” but how to represent and integrate geometry so that it contributes distinct signal beyond strong sequence priors. We provide architecture-agnostic diagnostics to test and quantify when and how explicit structure delivers that added value.

## Introduction

Predicting how single amino-acid substitutions (mutations) affect protein stability remains central to understanding disease mechanisms and engineering proteins for biotechnology. [1] Among stability-related phenotypes, changes in melting temperature (Δ*T_m_*) are particularly challenging: they reflect fold-level thermodynamics governed by distributed packing interactions, long-range contacts, and energetic perturbations traditionally thought least reducible to local sequence patterns. [2, 3]

Protein language models (PLMs) trained on vast protein sequence databases produce contextual residue representations encoding evolutionary constraints and biochemical regularities at unprecedented scale. [4–7] Yet the prevailing architectural paradigm for protein prediction tasks where knowledge suggest three-dimensional (3D) protein structure is important combines PLM embeddings with explicit 3D structure and auxiliary biochemical features, reflecting a widespread intuition: PLMs lack direct geometric supervision and may fail to capture how mutations propagate through protein structure. [4, 8] This intuition is most compelling for stability prediction. If any mutation-effect prediction task should benefit from explicit structural context, it ought to be Δ*T_m_*. Table 1 situates Δ*T_m_* among tasks traditionally regarded as highly structure-dependent. [9–14]

**Table 1:** Conceptual mapping between task type and expected dependence on explicit 3D structure.

| Task | Expected Structural Dependence | Notes |
| --- | --- | --- |
| **Fitness / growth / viability** | Low–Medium | Often dominated by conservation and mutational tolerance; structural effects are indirect or averaged out in bulk assays. |
| **Functional effects**<br>(activity, binding) | Medium | Active-site geometry and interfaces matter, but many effects are local; PLMs may capture a large fraction of signal. |
| **Pathogenicity** | Medium | Combines biochemical effects and stability with biological context; many predictors remain largely sequence-driven. |
| **$\Delta\Delta G$ (stability change)** | High | Directly tied to packing and long-range contacts; explicit 3D context is expected to be informative. |
| **$\Delta T_m$ (thermal stability shift)** | High | Reflects folding/unfolding thermodynamics; distributed structural perturbations are expected to matter. |

### The Challenge: Does Structure Add Value beyond PLM-derived Sequence Embeddings?

Despite the proliferation of multimodal “sequence + structure” models, a fundamental question is currently difficult to answer: *how much does explicit structure contribute beyond what PLMs already encode, and when does it contribute?* [15–17] Most studies introduce structural channels alongside architectural changes, different training data, or modified objectives, making it non-trivial to isolate structure’s marginal value. This attribution gap is increasingly consequential: structure prediction and geometric feature extraction impose substantial computational cost, yet if geometry provides limited incremental benefit, the default inclusion of expensive structural pipelines becomes hard to justify. [9, 10, 18–20]

### A Critical Problem: Structural Representations May Erase Mutation Effects

#### Key Insight: Representational Attenuation

**Mutation-conditioned structural information is not automatically present just because a model has a “structure channel.”**

Modern structure predictors (AlphaFold, ESMFold, RoseTTAFold) are trained to predict native structures from sequences using multiple sequence alignment (MSA) information. This means they inherently account for natural sequence variability, **effectively erasing the impact of mutation(s).** [18–20]

Moreover, standard graph constructions extracted from predicted structures, often the modality through which structure is integrated in multi-modal models, can attenuate or entirely remove the degrees of freedom most affected by substitutions (e.g., side-chain repacking, local conformational changes). Even when computational tools generate “mutant structures,” downstream featurization at coarse C*_α_*-only (or even backbone-only) resolution may not capture potentially-important local perturbations.

**A model may appear “structure-aware” while receiving geometric inputs that are effectively variant***^a^***-invariant.** The burden of proof is on the model developer to demonstrate that mutation-conditioned signal is present in the consumed features, not merely that a “structure channel” exists architecturally.

*^a^*Variant is a general term for a protein whose amino-acid sequence differs from the most common, or “wild-type,” sequence, typically resulting from a change (one or more mutations) in the corresponding gene

### Two Distinct Mechanisms: Static Context vs. Mutation-Induced Changes

In this paper, we distinguish two fundamentally different ways in which structure could help:

- **H1 (Static fold prior)**: Wild-type geometry provides environmental context (packing density, solvent exposure, neighborhood composition) that helps interpret mutations even when the structural representation is effectively identical for wild-type and variant inputs.
- **H2 (Mutation-conditioned perturbation)**: Differences between wild-type and variant geometries encode how substitutions alter contacts and stereochemistry, providing direct mechanistic signal about structural consequences.

These mechanisms have different scientific meanings and practical costs. H2 requires computationally generating variant structures and ensuring representations preserve geometric perturbations, both non-trivial. Yet they are rarely distinguished in literature despite having different implications for model design and resource allocation.

### Our Approach: Controlled Experiments to Isolate the Role of Tertiary Structure

We develop a controlled multimodal framework for Δ*T_m_* prediction to answer three questions:

1. **Which mechanism operates?** Do models benefit from mutation-conditioned structural perturbations (H2), or does a static wild-type fold prior (H1) suffice?
2. **Which inputs matter?** We perform systematic channel ablations holding model architecture and training constant, quantifying each input family’s marginal contribution.
3. **Where is the predictive signal located?** We introduce *context-radius probing* : training models restricted to progressively larger neighborhoods around mutations to test whether long-range context is actually used.

We complement aggregate performance metrics with variant-level diagnostics to characterize *when* additional modalities help.

### Key Findings

Across ten independent training runs per condition, our findings challenge conventional assumptions:

- **Sequence embeddings dominate.** Removing structure and auxiliary channels causes minimal performance change, while removing PLMs causes performance collapse. A PLM-only model matches the full multimodal performance.
- **Signal is highly local.** Mutation-site-only models recover full-context performance. Small windows (*r* ≤ 10 residues) saturate accuracy; most learnable signal is already in local PLM representations.
- **Static structure suffices; mutation-conditioned changes add little.** Providing the same wild-type structure for both (wild-type and variant) branches (H1) matches performance when (wild-type and variant) structures reportedly differ (H2). Geometric perturbations are not exploited to improve generalization.
- **Structure helps selectively, not uniformly.** Benefits concentrate in variants with atypical PLM embeddings occupying phenotypically-incoherent neighborhoods, regions where sequence-derived priors are locally unreliable.

These results do not imply structure is irrelevant to thermal stability. Rather, they reveal a **representation and integration gap**: current structural features and (modality) integration strategies fail to translate geometric information into robust gains beyond PLM priors, except potentially as sparse corrections for representational outliers.

The central challenge in multimodal variant-effect prediction is not *whether structure matters in principle*, but *whether we can*: (i) Provide mutation-sensitive structural representations that preserve perturbation signal; (ii) Integrate them selectively when PLM priors are unreliable; and (iii) Design architectures that exploit long-range couplings rather than redundant local context.

## 1 Related Work

### Structure-based stability prediction

For phenotypes coupled to physical energetics, particularly ΔΔ*G* and Δ*T_m_*, explicit 3D structure has been the primary predictive signal. Classical approaches estimate mutation impact (also known as variant effect) from local environments using statistical potentials (PoPMuSiC), energy functions (FoldX), or learned models (AUTO-MUTE, HoTMuSiC). [11–14, 21] Graph-based encodings (mCSM family) and 3D convolutional architectures (ThermoNet) extend this paradigm with deep learning. [22–26] This literature is tightly coupled to curated thermodynamic databases (ProTherm/ProThermDB, FireProtDB) providing the empirical foundation for expecting structure to matter for Δ*T_m_*. [27–31]

### From evolutionary features to protein language models

Sequence-only prediction progressed from conservation profiles to coevolutionary models (EVmutation) and deep generative models (DeepSequence, EVMutation). [32–34] Protein language models (PLMs) trained on massive unlabeled corpora (ESM-2, ProtTrans, ProteinBERT) now provide contextual residue representations encoding biochemical constraints and evolutionary regularities at scale. [4–7, 35] PLMs can be used zero-shot (masked-marginal likelihood) or as supervised features. [36, 37] Critically, PLMs often recover signals correlated with structural organization despite lacking explicit geometric supervision, making them competitive even for biophysical phenotypes. [1, 8] This shifts the burden of proof: multimodal designs must *demonstrate* structure adds non-redundant signal beyond modern sequence representations.

### Multimodal sequence and structure architectures

Recent predictors augment PLM embeddings with structural representations from experimental structures or AlphaFold/RoseTTAFold/ESM-Fold predictions. [18–20] Common architectures pair residue-level PLM features with geometric encoders, often graph neural networks operating on coordinate-derived graphs. Examples include GVP-GNN backbones, equivariant architectures (SE(3)-Transformers, E(n)-GNNs), and recent predictors combining PLM embeddings with structure-based encoders for pathogenicity, stability, and binding. [15, 16, 38–40] However, the statement that “structure helps” remains underspecified: does structure contribute as (i) a *static fold prior* providing wild-type environmental context, or (ii) *mutation-conditioned perturbation* exploiting geometric differences between wild-type and mutant structures? This distinction is rarely made explicit despite having different scientific and practical implications.

### The variant structure challenge and representational attenuation

Experimentally-resolved structures of variants are scarce at benchmark scale. For this reason, multimodal pipelines rely on computational surrogates: structure prediction with potentially further side-chain repacking/refinement. **Critically, modern structure predictors, such as AlphaFold, are trained to predict native structures from sequences using MSA information, inherently accounting for natural variability and potentially erasing variant effects.** [18, 20] Even when structures that potentially capture variant effects are generated, typical downstream featurization attenuates or erases mutation-affected degrees of freedom. Standard C*_α_*-based graph constructions are insensitive to side-chain repacking and local conformational changes. This creates a failure mode where models are nominally “structure-aware” while receiving effectively variant-invariant representations. Improvements may reflect wild-type priors rather than exploitation of mutation-conditioned changes.

### The attribution gap

Architectural and training changes are frequently bundled with structural channels (different encoders, pretraining regimes, data splits), making gains difficult to attribute specifically to geometric information. [41] For stability, prior ΔΔ*G* evaluations emphasize that average improvements mask failure modes and biases, motivating diagnostic analyses. [9, 10] Despite expanded datasets from deep mutational scanning (DMS), [42, 43] few controlled studies for Δ*T_m_* isolate structure’s *marginal* contribution while holding learning setup, supervision, and training protocol fixed. This gap is consequential: structural dependence expectations are strongest for Δ*T_m_*, yet structural preprocessing costs are non-trivial.

### Our contribution

To begin to answer questions on the potentially additive role of structure, in this paper we anchor our study in a representative, state-of-the-art (SOTA) multimodal framework:

PLM embeddings plus geometric representations and lightweight covariates, integrated via geometric deep networks. [15–17, 38] We introduce two key methodological tools: *wild-type-shared versus mutation-conditioned structural regimes* isolating mutant geometry’s role, and *context-radius probing* systematically restricting accessible information to neighborhoods around mutations. Together with controlled interventions and variant-level diagnostics, these tools provide an explicit, testable framing of what it means for structure to “help.”

## Materials and Methods

### Study Design and Hypotheses

Our study addresses a fundamental ambiguity in multimodal mutation-effect prediction: when authors report that *“structure helps”*, what mechanism is responsible? We formalize two competing hypotheses:

#### H1 (Static fold prior)

Explicit geometry helps by providing wild-type structural context (packing, solvent exposure, neighborhood composition). Under H1, the structural representation may be effectively constant across variants, and improvements do not require mutation-conditioned geometric changes.

#### H2 (Mutation-conditioned perturbation)

Explicit geometry helps by providing differences between wild-type and variant structures that encode mutation-induced perturbations. Under H2, the model requires mutation-sensitive geometries and representations preserving those differences.

To test these hypotheses, we hold the learning problem, model class, and training protocol fixed while systematically varying: (i) which input channels are available, (ii) whether structural features are wild-type-shared or mutation-conditioned, and (iii) the accessible context via context-radius probing. All experiments use ten independent training runs with matched random seeds for fair comparison.

### Dataset and Task

We use the standard Δ*T_m_* benchmark split: S4346 (4,346 variants, multiple proteins) for training and S571 (571 single-site substitutions, 37 proteins) for evaluation. [17] S571 spans diverse protein families, mutation types, and Δ*T_m_* magnitudes (range: −45.3 to +36.7 °C). The prediction task is as follows: given a wild-type protein *P_wt_* and single-residue substitution at position *i* (*a* → *b*), predict Δ*T_m_* = *T_m_*(*P_mut_*) − *T_m_*(*P_wt_*).

#### Potential inherent dataset biases

These datasets derive from curated stability databases (FireProtDB [31], ProTherm [28]) with known selection biases. Experimentally-characterized variants are typically chosen because researchers suspect measurable effects, often targeting functionally-important sites. This creates potential enrichment for (i) mutations with large local impact (visible to PLMs through sequence conservation), (ii) well-studied proteins, and (iii) structured regions rather than disordered loops. We note that this bias may partially explain our finding that mutation-site information is nearly sufficient: if variants predominantly affect local environments, long-range context naturally becomes less critical. However, this bias *strengthens* rather than weakens our core argument: even for thermal stability where distributed structural effects matter mechanistically, and even on variants likely enriched for structural impact, explicit geometry provides limited global benefit beyond PLM representations. Comprehensive deep mutational scanning datasets with neutral and deleterious mutations spanning full sequence space would provide valuable tests of whether structural contributions increase where PLM priors are weaker by design.

### Multimodal Graph Attention Framework (MMGAT)

In this paper we focus on a SOTA multimodal (MM) architecture template for stability prediction that integrates multiple residue-level representations via a geometric deep learning backbone [17]. This template, which we refer to as the *Multimodal Graph Attention Transformer* (MMGAT) from now on, serves as a unifying framework for testing different combinations of structural and sequence-derived modalities. A structure in this framework is represented as a graph whose nodes correspond to residues and whose edges reflect inter-residue geometric relationships. Residue-level inputs include features from three sources: (i) PLM-derived sequence embeddings, (ii) structure-derived pairwise and per-residue features, and (iii) auxiliary covariates, such as physicochemical descriptors and assay metadata. Each feature family is processed through dedicated input pathways and integrated within a GAT encoder that performs message-passing over the residue graph.

As in [17], we instantiate this template with six feature families per branch: (1) pairwise geometric descriptors (an *L* × *L* × 7 tensor encoding rotation/translation-invariant inter-residue geometry), (2) atom-presence masks, (3) per-residue structure confidence (pLDDT), (4) physicochemical amino-acid descriptors, (5) experimental assay pH, and (6) token embeddings from ESM-2 [4] (dimension 1280). Structure-derived inputs are varied in our study so as to investigate the previously-articulated hypothesis on what and where structure helps, if at all. In particular, in our study we include instantiations of this framework (ablations) where structure-derived inputs are computed from ESMFold-predicted 3D structures for both wild-type and variants. The two branches are processed independently and then integrated via a joint head to predict scalar Δ*T_m_*.

This MMGAT framework subsumes many recent multimodal designs [44–49] in mutation effect prediction and related tasks, including the GeoStab suite [17], whose GeoΔ*T_m_* model represents a specific instantiation of MMGAT for Δ*T_m_* prediction using a GAT-based encoder and all available input channels. We refer to this full-channel configuration as MMGAT^*^. Rather than treating GeoΔ*T_m_* as a fixed model under evaluation, we treat MMGAT as a design space and analyze its behavior through controlled interventions over input modality availability, structure regime, and context scope.

### Are Structural Features Mutation-conditioned?

We start with a question. Do geometric features consumed by the model contain mutation-specific information? The presence of a “structure channel” does not guarantee mutation-conditioned signal.

#### Released features are variant-invariant

We analyzed publicly-released S571 geometric tensors (GeoDTm-3D setting [17]) by comparing wild-type and variant representations. It is worth noting that in GeoDTm-3D, the wild-type structure is obtained from the Protein Data Bank (PDB) [50, 51], and the variant structure is modeled with FoldX [11] over the wild-type structure (with the mutation introduced to the sequence). The geometric tensors extracted from the wild-type and variant structures are analyzed and revealed to be nearly identical. Translation differences were *exactly zero*; orientation differences at numerical precision (∼ 10^−4^ radians). The geometric representation supplied to the model is effectively identical between wild-type and variant inputs despite separate channels. This establishes that in this regime, any benefit is necessarily consistent with **H1** (static fold prior) and not **H2**, because mutation-conditioned perturbations are not present. This is not surprisingly. FoldX can only model local structural perturbations, which are erased when the 3D structure is distilled into a residue interaction graph, which our analysis confirms.

While we do not have access to the geometric tensors supplied to the MMGAT framework in the GeoDTm-Seq regime in [17], we expect similar variant-invariance; in GeoDTm-Seq, wild-type structures are reported to be obtained via AlphaFold (though we note there is no reason to do so when experimentally-resolved structures exist for wild-type proteins), and variant structures are then obtained with FoldX-modifications over the AlphaFold wild-type structures to potentially account for the amino-acid mutation in a variant. As noted earlier, the interaction graph removes all potential local structural changes that FoldX may capture in a variant; the retained geometric descriptors do not encode side-chain geometry, so repacking is not retained.

#### Mutation-conditioned geometry is representable when variant structures are generated independently

Instead, in this paper, we introduce another regime where structures are generated separately for wild-type and variant sequences using ESMFold. We observe non-zero geometric differences (mean C*_α_* distance change 0.138Å). Critically, this is 3–4 times larger than ESMFold run-to-run variability (0.045Å), suggesting measurable differences in the wild-type versus variant channels that feed structural information to the GAT framework.

#### Experimental design: two structural regimes

These analyses motivate comparing two regimes:

- **Mutation-conditioned (testing H2)**: Generate ESMFold structures separately for a wild-type and variant pair. Compute all structure-derived features independently. Geometric tensors contain mutation-specific information (verified: mean |Δ*d*| ≈ 0.138 Å).
- **Wild-type-shared (testing H1)**: Generate structure only for the wild-type. Provide identical wild-type geometric features to both branches of a wild-type variant pair. Only PLM sequence-derived features differ.

If mutation-conditioned geometry provides value beyond wild-type context, the mutation-conditioned regime should substantially outperform the wild-type-shared one. Comparable performance would indicate H1 suffices.

### Controlled Channel Interventions

Our design systematically varies input availability: (i) structural regime (wild-type-shared vs. mutation-conditioned), (ii) modality availability (single-family ablations), and (iii) context scope (radius probing, described in detail later in this section). This isolates each information source’s marginal contribution while holding the architecture and optimization constant.

Across all experiments, we adopt the MMGAT architecture described above. Structural inputs are derived from ESMFold-predicted wild-type and variant structures and provided as separate channels to the encoder. We remind the reader that we refer to the configuration in which all input channels are present as MMGAT^*^.

#### Modality ablations

To quantify each channel’s contribution, we perform single-family ablations: one input family replaced by an all-ones tensor of identical shape. This removes information without changing dimensionality or requiring architecture modifications. We ablate each of six feature families individually, plus a **PLM-only** condition where all non-PLM inputs were masked simultaneously.

### Context-Radius Probing

To assess the locality of predictive signal and test whether long-range context is required for accurate prediction, we conduct a systematic radius-based intervention. For a mutation at residue *i*, we restrict each input feature tensor to a symmetric window of residues [*i* − *r, i* + *r*], where *r* ∈ {0, 1, 2*, . . . ,* 10, 20, 40*, . . . ,* 220}. A radius of *r* = 0 corresponds to a mutation-site-only context, while large radii approach the full sequence. All feature families—including PLM embeddings, geometric descriptors, and auxiliary features—respect the same radius mask.

Context-radius probing serves dual purposes: (i) diagnostic, revealing the spatial distribution of predictive signal (local vs. long-range), and (ii) design intervention, quantifying the minimum context required for accurate prediction. While implemented here within the MMGAT framework, this method is architecture-agnostic and so can be adapted to other architectures by applying the same radius-masking strategy across input channels.

### Training Protocol and Reproducibility

All models use identical loss functions: MSE plus a differentiable rank-based term encouraging Spearman correlation agreement (soft-rank loss, *λ* = 0.1). [52] Models are optimized using Adam (lr=10^−3^) for up to 50 epochs with early stopping (patience 10) based on validation Spearman correlation.

Each condition is trained for **ten independent runs** with distinct random seeds. Conditions include: full-input model (both structural regimes), each ablation, PLM-only model, and each context radius (23 radii × 10 runs = 230 models). Critically, **the same ten seeds are reused across all conditions**, so performance differences reflect input availability rather than initialization. Training/validation use protein-disjoint splits ( 10% validation). The test set (S571) is held separate throughout.

This design enables direct statistical comparison: for any two conditions, we have ten paired observations (same seed, different inputs), allowing paired significance tests.

### Baselines and Evaluation

We compare against: (i) **zero-shot PLM baseline** (ESM-2 masked-marginal scores), (ii) **released pretrained Geo**Δ*T_m_* **model** [17] (not directly comparable due to DMS pretraining but provides context for absolute performance), and (iii) competitive **structure-based predictors** in literature (AUTO-MUTE, [13] HoTMuSiC [14]). The primary performance metric is **Spearman rank correlation (***ρ*), consistent with benchmark protocols. We additionally report Pearson correlation (*r*), RMSE, and *R*^2^. For each condition, we report distributions across ten runs and test differences using paired Wilcoxon signed-rank tests (*α* = 0.05).

### Variant-level Diagnostic Analyses

To characterize *when* explicit structure helps, we perform variant-level analyses comparing full and PLM-only models.

#### Error difference analysis

For each variant *v*, we compute absolute errors *ɛ*_full_(*v*) and *ɛ*_PLM_(*v*), and their difference Δ*ɛ*(*v*) = *ɛ*_full_(*v*) − *ɛ*_PLM_(*v*). Negative values indicate structure helps (full model more accurate); positive values indicate PLM-only better. Predictions averaged across ten runs for robustness.

#### Identifying predictors of when structure helps

We define threshold-dependent binary classification: for threshold *t* ≥ 0, variants with |Δ*ɛ*(*v*)| < *t* excluded (choice doesn’t matter); variants with Δ*ɛ*(*v*) ≤ −*t* labeled “structure-helped”; Δ*ɛ*(*v*) ≥ +*t* labeled “structure-hurt.” We evaluate candidate scalar predictors using AUROC and AUPRC across thresholds *t* ∈ [0, 3] °C. Candidates include: embedding distance to centroid, relative solvent accessibility (RSA), packing density, mutation-site pLDDT, |Δ*T_m_*|, ESM-2 masked-marginal score, and PLM-only ensemble disagreement.

#### Phenotype coherence in PLM embedding space

For each test variant *v* with mutation-site ESM-2 embedding *e*(*v*) ∈ ℛ^1280^, let *N_k_*(*v*) be its *k* nearest neighbors. We quantify *phenotype coherence* using dispersion of ground-truth labels:

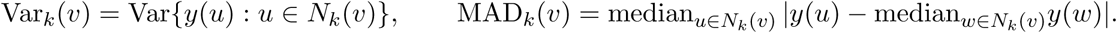

High Var*_k_* or MAD*_k_* indicates phenotype-incoherent regions where local proximity in representation space does not imply similar Δ*T_m_*. We report *k* = 10 (representative of observed robustness to *k* ∈ {15, 20, 25}).

### Implementation Notes

We reimplemented preprocessing and training from the published description [17] as complete training artifacts are not released. We did not use symmetry/inversion augmentation; all results correspond to non-augmented setting. We omitted the optional DMS pretraining phase because (i) full pretraining features are not released, (ii) regenerating structure-derived inputs at DMS scale is computationally prohibitive, and (iii) incorporating pretraining would require repeating it for each structural regime and modality condition, confounding our aim of isolating marginal contributions under controlled comparisons.

## 2 Results

We report results on the S571 Δ*T_m_* test set (571 variants, 37 proteins) using Spearman correlation (*ρ*) as primary metric, alongside RMSE and *R*^2^. Each condition is trained for ten independent runs with matched random seeds and protein-disjoint validation splits, so differences reflect information availability rather than initialization. Our results test two hypotheses: **H1** (structure as static fold prior) versus **H2** (structure via mutation-conditioned perturbations). We evaluate: (i) whether structural features actually differ between variants, (ii) global channel utility under controlled ablations, (iii) context requirements via radius probing, and (iv) regimes where geometry provides selective benefit.

### Do Structural Features Actually Change between Variants?

The presence of a “structure channel” does not guarantee the model receives mutation-specific information. We directly measured variant-sensitivity in the geometric representations consumed by the model.

#### Released features are variant-invariant

Analyzing publicly-released S571 geometric tensors (GeoΔ*T_m_* “3D” setting), we find translation differences exactly zero and orientation deviations at numerical precision (∼ 10^−4^ radians). The geometric representation is effectively identical between wild-type and variant input pairs. This establishes that any benefit in this regime is necessarily consistent with **H1** (static fold prior), not **H2** (mutation-conditioned perturbation), because mutation-conditioned perturbations are decidedly not present.

#### Mutation-conditioned signal is representable with independent structure generation

Generating structures separately for wild-type and variant sequences using ESMFold yields measurable wild-type variant differences (mean C*_α_* distance 0.138 Å), exceeding same-sequence ESMFold run-to-run variability (0.045 Å) under our settings. This indicates that the wild-type and variant channels are not trivially identical under independent structure generation, which is necessary for **H2** to be testable. We therefore evaluate two structural regimes: *wild-type-shared* (explicitly testing **H1**) and *mutation-conditioned* (making **H2** testable).

### Sequence Embeddings Dominate and Structure Adds Little Globally

Figure 1 provides a controlled attribution test: if a channel contributes non-redundant information, removing it should produce measurable, repeatable performance drops.

**Figure 1:**
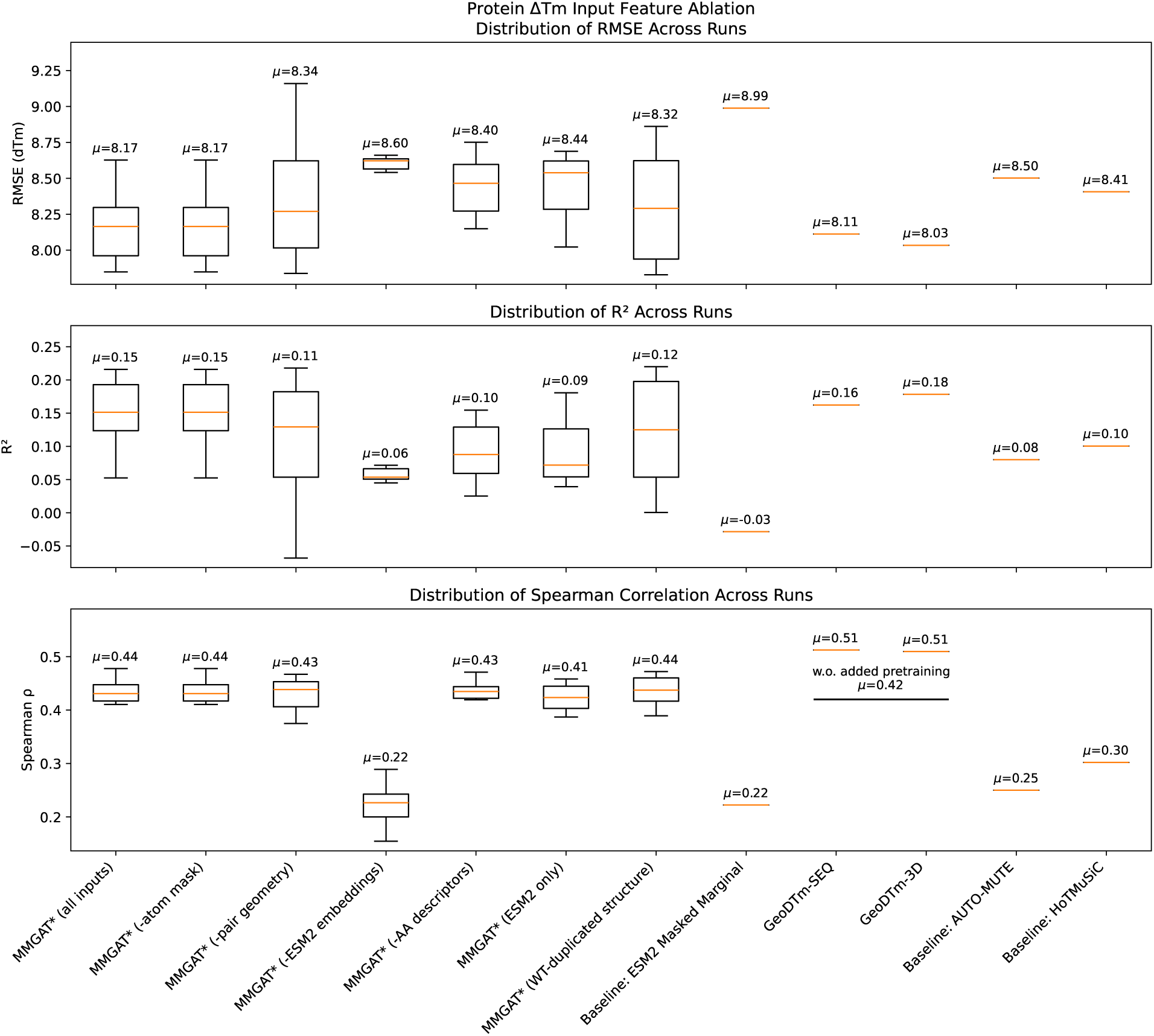
Global performance under controlled channel interventions. Performance across ten independent runs for full-input model, single-channel removals, and PLM-only condition. Mean values printed above boxes. External baselines included for context (not directly comparable when using additional pretraining or different structural regimes).

#### Removing PLM embeddings causes performance collapse

Ablating ESM-2 token embeddings yields the largest degradation (Spearman *ρ*: 0.44 → 0.22), reducing performance toward sequence-only baselines. PLMs provide a strong prior for mutation effects and are indispensable for competitive prediction in this setting.

#### Removing structure or auxiliary channels causes minimal change

In contrast, removing any single non-PLM family (geometric pair features, atom masks, structure confidence, physico-chemical descriptors, assay pH) produces marginal shifts relative to run-to-run variability. No single structural/auxiliary channel provides robust global improvement beyond PLM representations.

#### PLM-only model matches full multimodal performance

A model retaining only ESM-2 embeddings (all structure-derived and auxiliary inputs masked) overlaps the full-input distribution. Explicit structure and auxiliary covariates do not provide systematic global gains beyond PLM embeddings under this representation and training regime.

#### Scope

These findings are specific to the non-pretrained regime and current representations. They constrain what current multimodal designs extract from explicit geometry on this benchmark, not whether structure is irrelevant in principle.

### Static Fold Context Matches Mutation-Conditioned Geometry

We directly test whether mutation-conditioned structural perturbations provide value beyond a static wild-type fold prior by comparing two structural regimes under identical training (Figure 1, *MMGAT* (all inputs)* vs *MMGAT* (WT-duplicated structure)*).

#### Wild-type-shared geometry matches mutation-conditioned performance

Providing the same wild-type structure to both branches yields performance overlapping the mutation-conditioned regime across runs (mean *ρ*: 0.4365 vs 0.4361, paired Wilcoxon *p* = 0.77). Under current representations and integration, the model does not derive substantial global benefit from mutation-conditioned geometric differences beyond wild-type structural context. **H1** sufficiently explains the aggregate structural channel contribution.

#### Interpretation

The findings do not suggest that mutation-conditioned perturbations are absent in reality or never predictive. Rather, *given current structural representations and integration strategies*, mutation-conditioned perturbations are not translated into robust out-of-sample improvement on this benchmark. This is the integration challenge motivating the remaining analyses.

### Most Predictive Signal is Local, not Long-Range

We test whether the model uses broad sequence context via *context-radius probing* : training models restricted to symmetric windows [*i* − *r, i* + *r*] around mutation position *i* (Figure 2).

**Figure 2:**
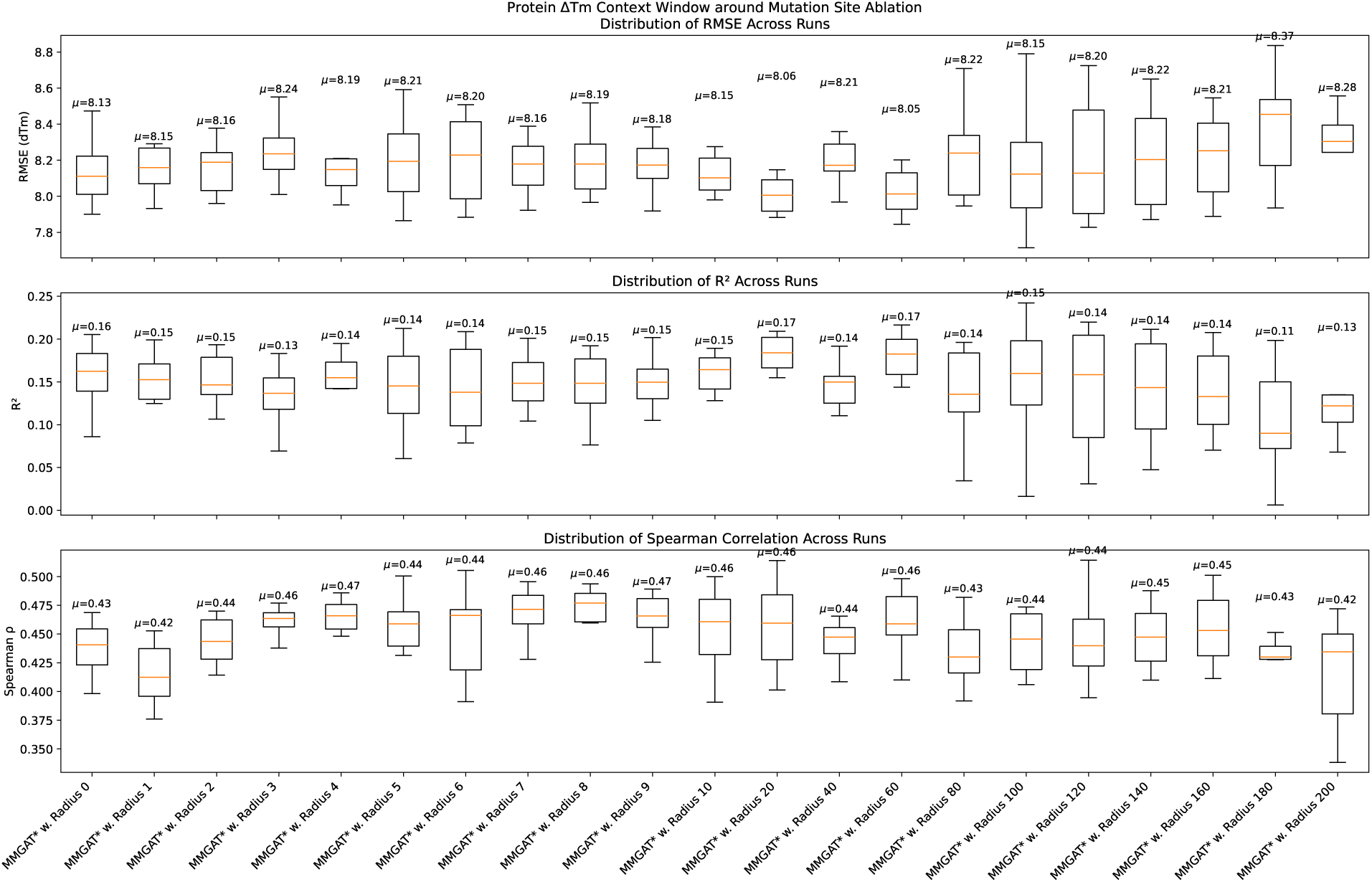
Context-radius dependence of Δ*T_m_* prediction. Performance across ten runs for models restricted to windows of radius *r* around mutations. Small windows match full-context accuracy; expanding to large radii yields limited gain.

#### Mutation-site information is nearly sufficient

Mutation-site-only models (*r* = 0) achieve performance close to full-context reference, indicating most learnable signal is in the mutation-site representation.

#### Modest local context saturates performance

Performance improves slightly at small radii then plateaus; expanding to global context provides no consistent out-of-sample gain. Even for Δ*T_m_*, traditionally regarded as long-range and structure-dependent, predictive signal exploited by this model class is overwhelmingly local.

#### Interpretation

This does not imply long-range interactions are unimportant for thermal stability mechanistically. Rather, under current representations and supervision, the model does not exploit long-range context in ways that improve generalization. This likely reflects both dataset bias toward variants with large local effects and limitations of current multimodal architectures in capturing distributed couplings.

### Embedding Distance Predicts Benefit, whereas Coarse Geometry Does Not

We define per-variant absolute errors and their difference Δ*ɛ* = *ɛ*_full_ − *ɛ*_PLM_. For threshold *t* ≥ 0, we label variants with Δ*ɛ* ≤ −*t* as structure-helped and Δ*ɛ* ≥ *t* as structure-hurt, then test which scalar signals discriminate these groups (Figure 3).

**Figure 3:**
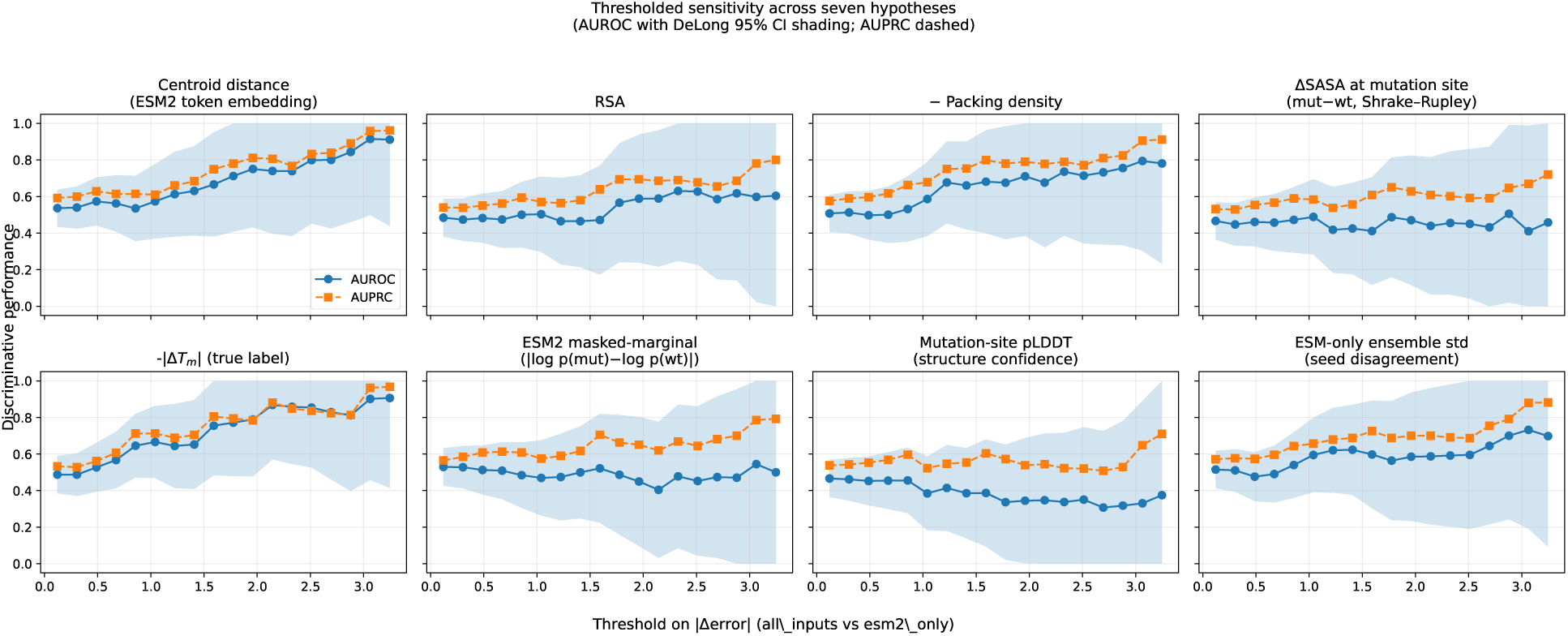
Predicting when explicit geometry helps. Each panel evaluates a candidate predictor. Solid curves: AUROC with 95% CI (DeLong); dashed: AUPRC. Strong predictors approach 1.0 as *t* increases (when model differences are large and reliable).

#### Distance to embedding centroid is the strongest signal

Across thresholds, **distance from mutation-site embedding centroid** most consistently identifies structure-helped variants, particularly at higher thresholds where model differences are large and less noise-dominated. The next two strongest predictors of structure helping, for high values of *t*, were very low values of ground-truth *δT_m_* and low packing density.

#### Coarse structural descriptors fail to discriminate

Relative solvent accessibility and other geometric environment measures do not consistently separate structure-helped from structure-hurt variants. Other candidate signals provide limited predictive value.

These results support a specific characterization: *explicit geometry helps where PLM representations are atypical*, not where simple geometric environment measures are extreme.

### Benefits Concentrate in Phenotypically-incoherent Neighborhoods

Distance-to-centroid suggests structure helps for representational outliers, but does not explain *why* those PLM representations are unreliable. We measure *phenotype coherence* of local neighborhoods using kNN dispersion of true Δ*T_m_* labels.

#### Structure-impact variants occupy label-incoherent regions

Variants where structure sub-stantially affects predictions show significantly higher neighborhood label dispersion (Figure 4). In these regions, local proximity in PLM space does not imply similar Δ*T_m_* values. The PLM has learned a representation misaligned with this phenotype locally, creating opportunity for orthogonal information (structure) to provide corrective signal. This pattern is robust across neighborhood sizes (*k* ∈ {10, 15, 20}).

**Figure 4:**
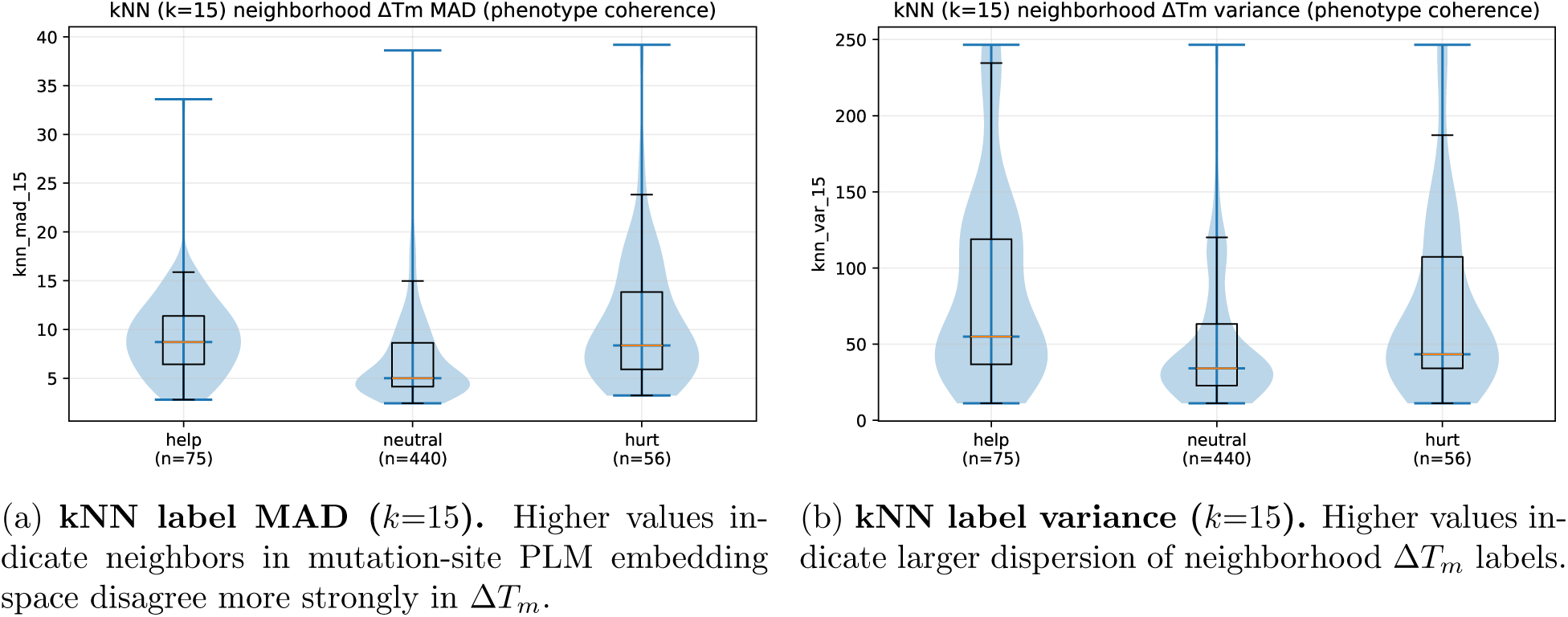
Phenotype-incoherent neighborhoods in mutation-site PLM embedding space. Violin plots show kNN-based phenotype coherence diagnostics across structure-helped, neutral, and structure-hurt variant groups. Structure-helped variants (Mann-Whitney U test vs neutral: MAD *p <* 0.001, Var *p <* 0.001) occupy neighborhoods where local proximity does not imply similar phenotype, consistent with locally unreliable sequence-derived priors.

### Summary of Findings

Across structural-regime tests, controlled ablations, context-radius probing, and variant-level diagnostics, we observe a consistent picture:

- **Variant-invariant representations are common:** At the level of consumed geometric tensors, “structure” can reduce to a wild-type fold prior when representations are effectively identical across variants. When variant-specific structures are generated independently, mutation-conditioned differences are present and exceed folding noise, making **H2** testable in principle.
- **PLM embeddings dominate globally:** Removing non-PLM channels yields limited change; removing PLM representations substantially degrades performance. A PLM-only model matches full multimodal performance.
- **Signal is highly localized:** Context-radius probing shows most learnable signal at mutation sites and nearby neighborhoods. Expanding to global context provides limited benefit, likely reflecting both dataset bias and architectural limitations.
- **Structure helps selectively:** Where explicit geometry improves predictions, benefits concentrate in variants atypical in PLM embedding space occupying phenotypically-incoherent neighborhoods. Coarse geometric descriptors provide limited explanatory power.

These findings support the paper’s framing: the central challenge is not whether structure matters in principle, but whether current representations and integration strategies provide and exploit mutation-sensitive geometric information yielding robust, generalizable gains beyond PLM priors.

## 3 Discussion and Conclusion

A central assumption in variant-effect prediction is that explicit 3D structure contributes essential information beyond what PLMs can infer from sequence. We test this assumption in a controlled setting for mutation-induced melting temperature shifts, Δ*T_m_*, a phenotype expected to depend strongly on protein geometry and stability energetics. We observe that PLM embeddings dominate performance, predictive signal is highly localized, and explicit geometry provides selective rather than uniform improvements.

### Representational attenuation and the H1/H2 distinction

Multimodal architectures can appear structure-aware while consuming effectively variant-invariant geometric features. We distinguish two mechanisms: H1 (static fold prior, where wild-type geometry provides environmental context) and H2 (mutation-conditioned perturbation, where geometric differences encode structural consequences). Our analyses show that current representations function primarily via H1. Even when variant structures are generated independently, standard residue-level featurizations (C*_α_*-based graphs, fixed connectivity) attenuate mutation-affected degrees of freedom.

### Why structure adds little for a structure-linked phenotype

At first glance, these results appear counterintuitive given that Δ*T_m_* reflects fold-level thermodynamics. The explanation lies in great part in a mismatch between how geometry determines stability and how geometry is represented in prevalent multimodal predictors. PLMs encode strong proxies for stability constraints through evolutionary statistics. Residue-level geometric descriptors from predicted structures may be too coarse, noisy, or misaligned with PLM-induced decision boundaries to provide robust incremental signal. Optimization preferentially exploits high-utility channels unless architectures or objectives explicitly incentivize weaker auxiliary inputs. The practical implication is consistent across explanations: naively adding structure does not ensure models use it to improve generalization. Recent work supports representation-aware diagnostics for when PLM-derived signals are reliable [53–55].

### Dataset considerations

The S4346/S571 benchmark inherits known selection biases from curated stability databases: variants are chosen for measurable effects and enriched at functionally important sites. This bias may contribute to observed locality if curated variants have strong local determinants already captured by PLM priors. However, this strengthens rather than weakens our empirical claim: even for a phenotype where structural dependence is widely presumed, and on variants likely enriched for structural impact, explicit geometry provides limited global benefit beyond PLMs under standard representations and integration strategies.

### Context-radius probing as diagnostic and design primitive

Beyond its role as a diagnostic for localizing predictive signal, the context-radius probing we propose in this paper additionally offers a practical design alternative when explicit 3D structure is unavailable or too costly to compute at scale: it enables controlled incorporation of progressively broader (sequence) context while keeping the learning problem and architecture fixed. The striking observation that mutation-site (or small-radius) models recover full-context performance in this benchmark therefore carries two implications: (i) the marginal utility of adding more context is empirically testable and often limited under current supervision and datasets, and (ii) when structure is omitted, carefully controlled sequence-context expansion can serve as a compute-light proxy for capturing any non-local effects the model is able to exploit.

### Why locality can coexist with “non-local” PLMs, and when structure should matter

That predictive signal saturates at the mutation site does not contradict the fact that PLMs use attention to integrate non-local context; rather, it suggests that for this task and dataset, the usable information for generalization is either already distilled into the local token representation or the model cannot reliably convert additional context into improved out-of-sample predictions. This immediately reframes the “when is structure interesting?” question: explicit structure should matter most precisely when sequence-derived priors are unreliable or uneven (e.g., in representational “junkyards” where embeddings are poorly calibrated [54], and in the twilight zone of low homology where PLM embeddings no longer reliably encode structural constraints [8]). Consistent with this view, our own variant-level analyses in this paper show that structure’s gains concentrate in atypical embedding regions with phenotypically-incoherent neighborhoods, indicating that geometry is most valuable as a targeted corrective signal rather than a uniformly beneficial add-on.

### Scope and future directions

We note that these conclusions are empirical and scoped to PLM-centered multimodal predictors using residue-level geometric descriptors on a standard Δ*T_m_* benchmark. They do not rule out settings where structure contributes larger gains: different supervision, all-atom or physics-informed representations, alternative pretraining regimes, or datasets enriched for cases where sequence statistics are insufficient. The central question is not whether structure matters mechanistically, but whether current representations and integration strategies allow it to contribute robustly (and where it matters) beyond PLM priors.

## Acknowledgments

This work was supported in part by the National Science Foundation Grant No. 2411529 and Grant No. 2310113. Computations were run on Hopper, a research computing cluster provided by the Office of Research Computing at George Mason University, VA (http://orc.gmu.edu).

## Notes

### Competing Interest Statement

The authors have declared no competing interest.

## References

[1] Yana Bromberg, R. Prabakaran, Anowarul Kabir, and Amarda Shehu. Variant effect prediction in the age of machine learning. Cold Spring Harbor Perspectives in Biology, 16(7):a041467, 2024. doi: 10.1101/cshperspect.a041467.

[2] Benjamin BV Louis and Luciano A Abriata. Reviewing challenges of predicting protein melting temperature change upon mutation through the full analysis of a highly detailed dataset with high-resolution structures. Molecular Biotechnology, 63(10):863–884, 2021.

[3] Lavi S Bigman and Yaakov Levy. Stability effects of protein mutations: the role of long-range contacts. The Journal of Physical Chemistry B, 122(49):11450–11459, 2018.

[4] Alexander Rives, Joshua Meier, Tom Sercu, et al. Biological structure and function emerge from scaling unsupervised learning to 250 million protein sequences. Proceedings of the National Academy of Sciences, 118(15):e2016239118, 2021. doi: 10.1073/pnas.2016239118.

[5] Ahmed Elnaggar, Michael Heinzinger, Christian Dallago, Ghalia Rihawi, Yu Wang, Llion Jones, Tom Gibbs, Tamas Feher, Christoph Angerer, Martin Steinegger, Debsindhu Bhowmik, and Burkhard Rost. Prottrans: Toward understanding the language of life through self-supervised learning. IEEE Transactions on Pattern Analysis and Machine Intelligence, 44(10):7112–7127, 2022. doi: 10.1109/TPAMI.2021.3095381.

[6] Roshan Rao, Nicholas Bhattacharya, Neil Thomas, Yan Duan, Xi Chen, John Canny, Pieter Abbeel, and Yun S. Song. Evaluating protein transfer learning with TAPE. In Advances in Neural Information Processing Systems (NeurIPS), volume 32, pages 9689–9701, 2019.

[7] Ethan C. Alley, Grigory Khimulya, Surojit Biswas, Mohammed AlQuraishi, and George M. Church. Unified rational protein engineering with sequence-based deep representation learning. Nature Methods, 16(12):1315–1322, 2019. doi: 10.1038/s41592-019-0598-1.

[8] Anowarul Kabir, Asher Moldwin, Yana Bromberg, and Amarda Shehu. In the twilight zone of protein sequence homology: do protein language models learn protein structure? Bioinformatics Advances, 4(1):vbae119, 2024. doi: 10.1093/bioadv/vbae119.

[9] Vladimir Potapov, Matan Cohen, and Gideon Schreiber. Assessing computational methods for predicting protein stability upon mutation: good on average but not in the details. Protein Engineering, Design & Selection, 22(9):553–560, 2009. doi: 10.1093/protein/gzp030.

[10] Grant Thiltgen and Richard A. Goldstein. Assessing predictors of changes in protein stability upon mutation using self-consistency. PLOS ONE, 7(10):e46084, 2012. doi: 10.1371/journal.pone.0046084.

[11] Raphael Guerois, Jens Erik Nielsen, and Luis Serrano. Predicting changes in the stability of proteins and protein complexes: a study of more than 1000 mutations. Journal of Molecular Biology, 320(2):369–387, 2002. doi: 10.1016/S0022-2836(02)00442-4.

[12] Yves Dehouck, Aline Grosfils, Benjamin Folch, Dimitri Gilis, Philippe Bogaerts, and Marianne Rooman. Fast and accurate predictions of protein stability changes upon mutations using statistical potentials and neural networks: PoPMuSiC-2.0. Bioinformatics, 25(19):2537–2543, 2009. doi: 10.1093/bioinformatics/btp445.

[13] Majid Masso and Iosif I. Vaisman. AUTO-MUTE: web-based tools for predicting stability changes in proteins due to single amino acid replacements. Protein Engineering, Design & Selection, 23(8):683–687, 2010. doi: 10.1093/protein/gzq042.

[14] Fabrizio Pucci, Raphaël Bourgeas, and Marianne Rooman. Predicting protein thermal stability changes upon point mutations using statistical potentials: Introducing HoTMuSiC. Scientific Reports, 6:23257, 2016. doi: 10.1038/srep23257.

[15] Zichen Wang, Steven A. Combs, et al. LM-GVP: an extensible sequence and structure informed deep learning framework for protein property prediction. Scientific Reports, 12:6832, 2022. doi: 10.1038/s41598-022-10775-y.

[16] Peng Cheng, Cong Mao, Jin Tang, Sen Yang, Yu Cheng, Wuke Wang, Qiuxi Gu, Wei Han, Hao Chen, Sihan Li, et al. Zero-shot prediction of mutation effects with multimodal deep representation learning guides protein engineering. Cell Research, 34(9):630–647, 2024.

[17] Yunxin Xu, Di Liu, and Haipeng Gong. Improving the prediction of protein stability changes upon mutations by geometric learning and a pre-training strategy. Nature Computational Science, 4(11):840–850, 2024.

[18] John Jumper, Richard Evans, Alexander Pritzel, et al. Highly accurate protein structure prediction with AlphaFold. Nature, 596:583–589, 2021. doi: 10.1038/s41586-021-03819-2.

[19] Minkyung Baek, Frank DiMaio, Ivan Anishchenko, et al. Accurate prediction of protein structures and interactions using a three-track neural network. Science, 373(6557):871–876, 2021. doi: 10.1126/science.abj8754.

[20] Zeming Lin, Halil Akin, Roshan Rao, et al. Evolutionary-scale prediction of atomic-level protein structure with a language model. Science, 379(6637):1123–1130, 2023. doi: 10.1126/science.ade2574.

[21] Joost Schymkowitz, Jesper Ferkinghoff-Borg, François Stricher, Robby Nys, Frederic Rousseau, and Luis Serrano. The FoldX web server: an online force field. Nucleic Acids Research, 33 (Web Server issue):W382–W388, 2005. doi: 10.1093/nar/gki387.

[22] Douglas E. V. Pires, David B. Ascher, and Tom L. Blundell. mCSM: predicting the effects of mutations in proteins using graph-based signatures. Bioinformatics, 30(3):335–342, 2014. doi: 10.1093/bioinformatics/btt691.

[23] Douglas E. V. Pires, David B. Ascher, and Tom L. Blundell. Duet: a server for predicting effects of mutations on protein stability using an integrated computational approach. Nucleic Acids Research, 42(W1):W314–W319, 2014. doi: 10.1093/nar/gku411.

[24] Carlos H. M. Rodrigues, Douglas E. V. Pires, and David B. Ascher. DynaMut: predicting the impact of mutations on protein conformation, flexibility and stability. Nucleic Acids Research, 46(W1):W350–W355, 2018. doi: 10.1093/nar/gky300.

[25] Hongjian Cao, Junjie Wang, Luhua He, Yao Qi, and Jianlin Z. H. Zhang. DeepDDG: Predicting the stability change of protein point mutations using neural networks. Journal of Chemical Information and Modeling, 59(4):1508–1514, 2019. doi: 10.1021/acs.jcim.8b00697.

[26] Geng Li, Ming Dong, Huabin Fan, et al. ThermoNet: a deep 3d convolutional neural network for predicting protein stability changes. PLOS Computational Biology, 16(11):e1008291, 2020. doi: 10.1371/journal.pcbi.1008291.

[27] Michael M. Gromiha, Jun An, Hiroshi Kono, Masahiro Oobatake, Haruki Uedaira, and Akinori Sarai. ProTherm: thermodynamic database for proteins and mutants. Nucleic Acids Research, 27(1):286–288, 1999. doi: 10.1093/nar/27.1.286.

[28] Michael M. Gromiha, Haruki Uedaira, Jun An, Samuel Selvaraj, Ponraj Prabakaran, and Akinori Sarai. ProTherm: thermodynamic database for proteins and mutants: developments in version 3.0. Nucleic Acids Research, 30(1):301–302, 2002. doi: 10.1093/nar/30.1.301.

[29] K. A. Bava, Michael M. Gromiha, Haruki Uedaira, Keiko Kitajima, and Akinori Sarai. ProTherm, version 4.0: thermodynamic database for proteins and mutants. Nucleic Acids Research, 32(Database issue):D120–D121, 2004. doi: 10.1093/nar/gkh082.

[30] Ranjeet Nikam, Abhishek Kulandaisamy, K. Harini, Deepak Sharma, Michael M. Gromiha, Akinori Sarai, M. K. Prakash, and Narayanaswamy Srinivasan. ProThermDB: thermodynamic database for proteins and mutants revisited after 15 years. Nucleic Acids Research, 49(D1): D420–D424, 2021. doi: 10.1093/nar/gkaa1035.

[31] Jiri Stourac, Tomas Martinek, Jan Bendl, Jan Brezovsky, Laura Priegue, Carlos Azevedo, Ricardo Honorato, David Bednar, Zbynek Prokop, and Jiri Damborsky. FireProtDB: a database of thermostable proteins and mutations. Nucleic Acids Research, 49(D1):D319–D324, 2021. doi: 10.1093/nar/gkaa981.

[32] Thomas A. Hopf, John B. Ingraham, Frank J. Poelwijk, Charlotta P. I. Schärfe, Michael Springer, Chris Sander, and Debora S. Marks. Mutation effects predicted from sequence co-variation. Nature Biotechnology, 35(2):128–135, 2017. doi: 10.1038/nbt.3769.

[33] Adam J. Riesselman, John B. Ingraham, and Debora S. Marks. Deep generative models of genetic variation capture the effects of mutations. Nature Methods, 15(10):816–822, 2018. doi: 10.1038/s41592-018-0138-4.

[34] Jonathan Frazer, Pascal Notin, Mafalda Dias, Andrea Gomez, Jungo Min, Kelly Brock, Yarin Gal, and Debora S. Marks. Disease variant prediction with deep generative models of evolutionary data. Nature, 599(7883):91–95, 2021. doi: 10.1038/s41586-021-04043-8.

[35] Nadav Brandes, Dan Ofer, Yam Peleg, Nadav Rappoport, and Michal Linial. ProteinBERT: a universal deep-learning model of protein sequence and function. Bioinformatics, 38(8):2102–2110, 2022. doi: 10.1093/bioinformatics/btac020.

[36] Joshua Meier, Roshan Rao, Robert Verkuil, Jason Liu, Tom Sercu, and Alexander Rives. Language models enable zero-shot prediction of the effects of mutations on protein function. In Advances in Neural Information Processing Systems (NeurIPS), 2021. URL https://proceedings.neurips.cc/paper/2021/hash/f51338d736f95dd42427296047067694-Abstract.html.

[37] Roshan M. Rao, Jason Liu, Robert Verkuil, Joshua Meier, John Canny, Pieter Abbeel, Tom Sercu, and Alexander Rives. MSA transformer. In Proceedings of the 38th International Conference on Machine Learning (ICML), 2021. URL https://proceedings.mlr.press/v139/ rao21a.html.

[38] Bowen Jing, Stephan Eismann, Patricia Suriana, Raphael J. Townshend, and Ron Dror. Learning from protein structure with geometric vector perceptrons. In International Conference on Learning Representations (ICLR), 2021. URL https://openreview.net/forum?id=1YLJDvSx6J4.

[39] Fabian B. Fuchs, Daniel E. Worrall, Volker Fischer, and Max Welling. SE(3)-transformers: 3d roto-translation equivariant attention networks. In Advances in Neural Information Processing Systems (NeurIPS), 2020. URL https://proceedings.neurips.cc/paper/2020/hash/15231a7ce4ba789d13b722cc5c955834-Abstract.html.

[40] Víctor Garcia Satorras, Emiel Hoogeboom, and Max Welling. E(n) equivariant graph neural networks. In Proceedings of the 38th International Conference on Machine Learning (ICML), volume 139 of *Proceedings of Machine Learning Research*, pages 9323–9332, 2021. URL https://proceedings.mlr.press/v139/satorras21a.html.

[41] Pascal Notin, Aaron W. Kollasch, Daniel Ritter, et al. ProteinGym: Large-scale benchmarks for protein fitness prediction and design. *bioRxiv*, 2023.doi: 10.1101/2023.12.07.570727.

[42] Douglas M. Fowler and Stanley Fields. Deep mutational scanning: a new style of protein science. Nature Methods, 11(8):801–807, 2014. doi: 10.1038/nmeth.3027.

[43] Kotaro Tsuboyama, Justas Dauparas, Jonathan Chen, et al. Mega-scale experimental analysis of protein folding stability in biology and design. Nature, 620(7973):434–444, 2023. doi: 10.1038/s41586-023-06328-6.

[44] Haifan Gong, Yumeng Zhang, Chenhe Dong, Yue Wang, Guanqi Chen, Bilin Liang, Haofeng Li, Lanxuan Liu, Jie Xu, and Guanbin Li. Unbiased curriculum learning enhanced global-local graph neural network for protein thermodynamic stability prediction. Bioinformatics, 39(10): btad589, 2023.

[45] Yunzhuo Zhou, Qisheng Pan, Douglas EV Pires, Carlos HM Rodrigues, and David B Ascher. Ddmut: predicting effects of mutations on protein stability using deep learning. Nucleic Acids Research, 51(W1):W122–W128, 2023.

[46] Yuan Zhang, Junsheng Deng, Mingyuan Dong, Jiafeng Wu, Qiuye Zhao, Xieping Gao, and Dapeng Xiong. Pilot: Deep siamese network with hybrid attention improves prediction of mutation impact on protein stability. Neural Networks, 188:107476, 2025.

[47] Kanchan Jha, Sriparna Saha, and Hiteshi Singh. Prediction of protein–protein interaction using graph neural networks. Scientific Reports, 12(1):8360, 2022.

[48] Ramisa Alam, Sazan Mahbub, and Md Shamsuzzoha Bayzid. Pair-egret: enhancing the prediction of protein–protein interaction sites through graph attention networks and protein language models. Bioinformatics, 40(10):btae588, 2024.

[49] Zhonghui Gu, Xiao Luo, Jiaxiao Chen, Minghua Deng, and Luhua Lai. Hierarchical graph transformer with contrastive learning for protein function prediction. Bioinformatics, 39(7): btad410, 2023.

[50] Helen M. Berman, John Westbrook, Zukang Feng, Gary Gilliland, T. N. Bhat, Helge Weissig, Ilya N. Shindyalov, and Philip E. Bourne. The protein data bank. Nucleic Acids Research, 28 (1):235–242, 2000. doi: 10.1093/nar/28.1.235.

[51] wwPDB consortium. Protein data bank: the single global archive for 3d macromolecular structure data. Nucleic Acids Research, 47(D1):D520–D528, 2019. doi: 10.1093/nar/gky949.

[52] Mathieu Blondel, Olivier Teboul, Quentin Berthet, and Josip Djolonga. Fast differentiable sorting and ranking. In International Conference on Machine Learning, pages 950–959. PMLR, 2020.

[53] C. Hou, D. Liu, A. Zafar, and Y. Shen. Understanding protein language model scaling on mutation effect prediction. *bioRxiv*, 2025. doi: 10.1101/2025.04.25.650688.

[54] R. Prabakaran and Y. Bromberg. Quantifying uncertainty in protein representations across models and task. *bioRxiv*, 2025. doi: 10.1101/2025.04.30.651545.

[55] Tobias Senoner, Ivan Koludarov, Joshua Guenther, Amarda Shehu, Burkhard Rost, and Yana Bromberg. Which plm to choose? *bioRxiv*, 2025. doi: 10.1101/2025.10.30.685515.

